# Cross-family and phage-specific gene requirements for Klebsiella infection revealed by scalable RB-TnSeq genetic screens

**DOI:** 10.64898/2026.03.14.711032

**Authors:** Marissa R. Gittrich, Courtney M. Sanderson, Cara M. Noel, Erica Babusci, Sumeyra C. Selbes, Madeline Svab, Isabella Murray, Collis Bousliman, Ami Fofana, Aghiad Daboul, Jonathan Leopold, Alessandra Gonçalves de Melo, Marion Urvoy, Sylvain Moineau, Vivek K. Mutalik, Matthew B. Sullivan

## Abstract

Bacteriophages are being cataloged at an accelerating pace and are recognized as key players in nutrient and energy cycling across ecosystems. Yet the bacterial genetic determinants that govern phage-host specificity and infection success remain poorly understood, particularly in clinically and ecologically important genera such as *Klebsiella* where prior receptor characterization has been almost entirely limited to capsulated strains. Here we used a randomly barcoded, genome-wide, loss-of-function transposon mutant library (RB-TnSeq) of *Klebsiella* sp. M5al, a naturally acapsular, nitrogen-fixing rhizobacterium, to generate the first systematic, cross-family map of phage receptor gene dependencies in *Klebsiella*. Challenging the library against 25 double-stranded DNA phages spanning five families in 213 parallel assays, we identified 42 bacterial genes associated with phage infection, of which 15 had no prior association with phage infection in any bacterial system. Disruption of surface receptor biosynthesis genes conferred cross-resistance across multiple phage families, while intracellular gene disruptions had predominantly phage-specific effects. Clonal validation of eight genes confirmed LPS outer core biosynthesis genes as primary receptor determinants alongside additional host factors spanning outer membrane transport, cofactor biosynthesis, and two-component signaling. Comparative analysis across all 25 phages revealed that phage genus rather than family is the stronger predictor of host gene dependency profiles, a finding with direct implications for the functional annotation of uncharacterized phage isolates and rational phage cocktail design. Together, these findings provide a community resource for linking phage genomic diversity to functional host interaction space in this ecologically and clinically important genus.

**Author Summary:** Bacteriophages, or phages, are viruses that infect bacteria and play central roles in shaping microbial communities across ecosystems. Despite their ecological importance, the bacterial genes that determine infection outcomes remain poorly understood. Using a scalable barcoded transposon sequencing (RB-TnSeq) approach, we mapped host gene requirements for infection by 25 diverse phages of *Klebsiella* sp. M5al, a soil-associated plant-growth-promoting bacterium. We identified 42 bacterial genes associated with phage infection, spanning surface receptors, transcriptional regulators, and metabolic and protein-folding pathways. Infection strategies broadly clustered by phage genus and family, while phage-specific differences arose primarily in intracellular processes such as transcription and protein folding. These findings reveal both conserved and phage-specific host interactions that define infection strategies and establish a scalable framework for linking phage genomic diversity to function, with practical implications for predicting phage host range, designing phage cocktails for therapy, and understanding phage-driven dynamics in natural microbial communities.

## Introduction

Phages, viruses that infect bacteria, have been increasingly recognized as key players in ecological and evolutionary processes across Earth’s ecosystems (1–9). In terrestrial environments, phages have been found to impact nutrient cycling (10,11), shape bacterial community composition (12–14), and impact plant health (15,16). Despite over a century of phage research and growing recognition of their ecological importance, modeling phage-host interactions remains challenging due to limited studies of these interactions compared to the abundance and diversity of phages (17–19). Even the basic question of “who infects whom” is often difficult to answer as (i) nearly genetically identical phages can exhibit drastically different host-range and infectivity at the bacterial strain level (20–25), and (ii) infectivity can change based on environmental conditions (25–27).

Mechanistically, some phage susceptibility differences can be explained by variations in bacterial receptors and in phage receptor binding proteins (20,21,28), presence or absence of host surface proteins (22), or phage defense elements like restriction-modification and CRISPR-Cas systems (23). However, these factors do not fully explain the substantial variability in susceptibility observed across strains and phages. In particular, the broader and strain-specific genetic determinants that modulate phage infection beyond well-characterized receptors and systems remain incompletely defined (24,25). In addition, there is a large discrepancy between the number of culturable phage-host model systems available for experimental testing (maybe hundreds to thousands) and the uncultivated genomic fragments known to exist (>15 million in IMG-VR) (29). These genomic fragments represent only a small subset of global phage numbers thought to exist planet-wide (∼10^31^) (30). Critically, even the best *in silico* host prediction approaches are limited to predicting phage hosts to the bacterial genus level for uncultivated phage groups (31,32), which represents a significant bottleneck in phage-driven microbiome design efforts (32). These knowledge gaps in mechanistic understanding and understudied taxonomic variation in phage-host interactions make accurately predicting phage infectivity at the strain level challenging.

Recent advances in genome-wide functional genomics offer a scalable path toward resolving these gaps in phage-host interaction biology. Genome-wide transposon mutagenesis enables systematic interrogation of bacterial genes non-essential for growth under defined conditions and has been applied to characterize host factors involved in phage infection (33–36). The combination of this approach with high-throughput randomly-barcoded transposon-site sequencing (RB-TnSeq), enables cost-effective, parallel fitness assessment of more than 100,000 transposon mutants in a single pooled experiment, quantifying the contribution of each disrupted gene to bacterial survival under a given selective pressure, including phage challenge (37–40). Bacterial genes required for successful phage infection are identified through the selective enrichment of the mutated strains after phage exposure. This enrichment can also be quantified as a fitness advantage for the mutant attributed to that gene. Unlike traditional single-gene knockouts, RB-TnSeq provides genome-scale resolution of key host factors involved in phage infection pathways in a single sequencing run, substantially accelerating the systematic characterization of host factors governing phage infection.

RB-TnSeq has already revealed novel insights into phage biology, even in well-studied systems like *Escherichia coli* and *Salmonella enterica* (39,40). These studies have uncovered previously unrecognized host genes involved in phage susceptibility and highlighted genes conferring cross-resistance to diverse phages, such as genes involved in surface structures, transcriptional regulation, and stress responses. In one notable case, RB-TnSeq in *E. coli* revealed that cyclic di-GMP regulates secretion of the glycan receptor for phage N4, overturning a thirty-year-old model of phage adsorption and explaining the broad host range of N4-like phages (39,41–43). Most recently, while this manuscript was under review, Moriniere et al. (2026) (44) extended RB-TnSeq approach to an unprecedented scale, conducting genome-wide genetic screens across 255 taxonomically diverse *E. coli* phages and assigning receptor specificity to 193 phages across 19 receptor classes. By integrating these data with comparative genomics and machine learning, the authors demonstrated that phage receptor identity can be predicted from genome sequence alone, illustrating the predictive power that large-scale RB-TnSeq datasets can unlock.

Despite its clinical prominence as a leading cause of antibiotic-resistant infections, *Klebsiella* remains poorly characterized at the level of phage receptor biology (45). Our understanding of *Klebsiella* phage-host interactions has been shaped almost entirely by studies of capsulated clinical strains, in which the capsular polysaccharide (CPS) serves as the dominant surface feature and principal phage receptor (33,46–49). Recently, efforts have begun to systematically catalogue *Klebsiella* phage diversity, including the *Klebsiella* Phage Collection (50), as well as our prior work characterizing 24 Klebsiella phages that are publicly available through Université Laval (51), reflecting growing recognition of the need for organized phage resources in this genus. However, decades of research have uncovered only a handful of capsule-independent phage receptors (33,46), and there has been no systematic study on experimentally mapping a diverse phage to different host factor determinants. This capsule-dominated view of *Klebsiella* phage receptor biology has failed to account for the genotype that emerges most frequently under phage selection pressure. Under phage predation, *K. pneumoniae* most frequently evolves resistance by shedding or modifying its capsule, with more than 97% of phage-resistant mutants carrying CPS biosynthesis mutations in controlled experiments (49). Capsule loss can also arise reversibly without any genetic mutation (52). These acapsular escape variants are no longer susceptible to capsule-targeting phages and instead become susceptible to phages that use secondary receptor structures, namely LPS outer core and outer membrane porins. Characterizing receptors in acapsular strains therefore directly reveals the phage infection landscape relevant to environmental *Klebsiella* populations, where capsule-independent host-phage interactions are likely to predominate.

To address this gap, we selected *Klebsiella* sp. M5al, a naturally non-capsulated, nitrogen-fixing soil bacterium relevant to nutrient cycling and plant root colonization (53–55), as the experimental host. We hypothesized that the absence of capsular structure would reveal the underlying LPS– and porin-dependent receptor landscape that is obscured by capsule signals in capsulated strain screens and can serve as a model system for systematically characterizing secondary or terminal receptors. As a demonstration, using this host, we assembled a collection of 25 phages spanning five families, capturing substantial genomic and morphological diversity. We challenged the *Klebsiella* sp. M5al RB-TnSeq library against all 25 phages in parallel, identified 42 bacterial genes associated with phage infection, and generated the first systematic, cross-family map of receptor gene dependencies across a diverse panel of *Klebsiella* phages.

## Results and Discussion

### Challenging the *Klebsiella* sp. M5al RB-TnSeq library against 25 phages

We challenged *Klebsiella* sp. M5al RB-TnSeq library against 25 phages (51) spanning five families, including the three most prevalent among *Klebsiella* phages, *Autographiviridae* (n=3), *Drexlerviridae* (n=2), *Straboviridae* (n=13)], as well as two recently described novel families Ca. *Mavericviridae* (n=3) and Ca. *Rivulsuviridae* (n=4) (**Fig. 1A**). RB-TnSeq experiments were conducted as separate, single phage challenges at three phage-to-bacteria ratios (Methods, multiplicity of infection – MOI 10, 1, and 0.1). Strain fitness was assessed from the log2 barcode abundance changes from the 4.5 hours post-infection relative to the initial library composition (“T0”) with individual strain fitness values averaged across all barcode abundances in a given gene to yield a gene-level fitness score (**Fig. 1B**). To minimize false positives, we restricted analyses to genes meeting three criteria: (1) a fitness score of at least 4, representing the log_₂_ fold-change in mutant abundance relative to the no-phage control; (2) a t-like statistic of at least 5, reflecting consistency of the fitness estimate across independent barcodes; and (3) an estimated standard error no greater than 2, indicating high confidence in the calculated value. These thresholds were applied as previously described (39). Loss-of-function mutants meeting these criteria and exhibiting increased fitness were interpreted as conferring partial or complete resistance to phage infection.

**Figure 1.**
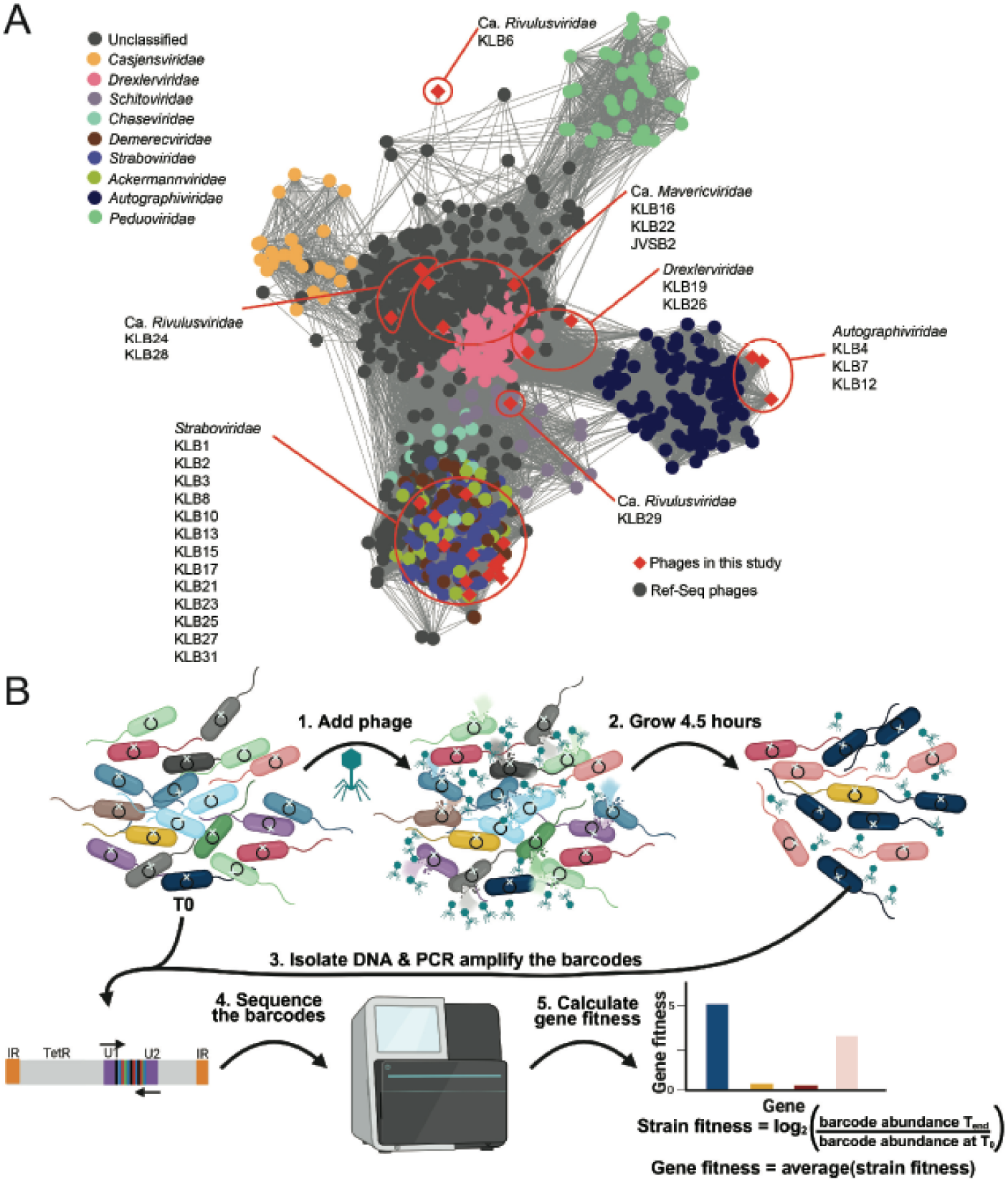
Overall study design. (A) Taxonomic diversity, determined through ‘vConTACT3’ gene sharing network analyses, for the 25 phages used in this study (red diamonds, and “KLB” named phages listed under each family name) compared to the 942 double-stranded DNA phages infecting *Klebsiella*, *Salmonella*, and *Escherichia* available from NCBI’s RefSeq (version 220, 1/10/24). Lines represent detectable gene content similarity between the phages, nodes represent phage genomes, and the phages are color-coded according to family (see legend). (B) Overview of the RB-TnSeq experiment. RB-TnSeq library is initially barcode-sequenced to establish starting mutant frequencies, then challenged with one of the 25 phages, incubated in a plate reader, and the resultant mutant frequencies are estimated through barcode-sequencing. For each transposon knockout, the strain-level fitness is calculated by comparing the barcode abundance at the end of the infection to the initial abundance at T0 (see equation in the figure). At least three transposon knockouts or strains were required to establish each gene’s average fitness.

This experimental and analytical approach was then used to explore the bacterial genes required for phage infection across 213 genome-wide assays, identifying 42 bacterial genes, with each phage required between 1 and 15 (median = 3) genes. The 42 genes span nine functional categories such as surface structures, inner membrane proteins, transcriptional regulators, carbon metabolism, energy cycling, amino acid biosynthesis and transport, protein transport, protein modification, and hypothetical proteins (**Fig. 2**). Of these, 25 genes encode surface-associated factors and are likely involved in phage adsorption, while the remaining 17 encode intracellular functions. All 25 phages produced at least one high-scoring receptor hit, and eight had hits for more than one putative receptor (**Fig. 2**).

**Figure 2.**
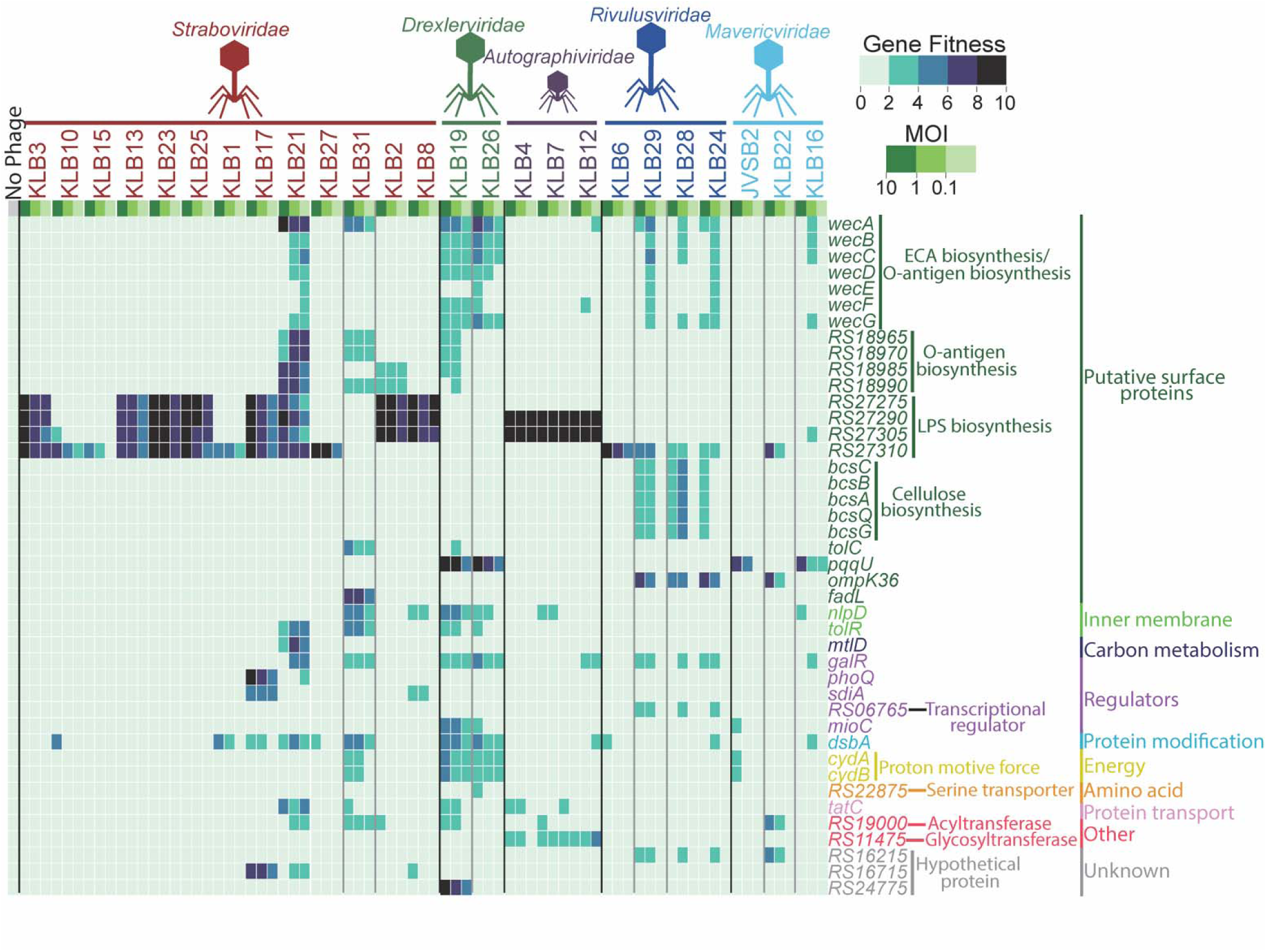
An RB-TnSeq mutant pool’s gene fitness against diverse phage infections. For each assay, a subset of a *Klebsiella* sp. M5al RB-TnSeq mutant pool was infected by a single phage (25 phages in total) at a multiplicity of infection (MOI) of 0.1, 1, or 10 (Methods). We also included ‘no-phage’ culture where the SM buffer was mixed with the library in place of phage. Gene fitness was then calculated as described above (**Fig. 1B**, showing high-confidence bacterial genes (a gene fitness score of ≥4). Each experiment was done in duplicate, and each gene had >3 unique gene knockouts. Vertical lines are used to separate phage families (black lines) or phage genera (gray lines). The underlying data for this figure can be found in **Table S1-S2**.

### Connecting phage taxonomy to host gene dependency profiles

The scale of this genome-wide screen, encompassing 25 phages across five families, enabled us to ask a question that single phage-host studies cannot address. Specifically, whether infection strategies are conserved within phage taxonomic groups or diverge even among closely related phages. To address this, we examined the extent to which RB-TnSeq fitness profiles cluster by phage taxonomy at the family and genus levels. Visual inspection revealed clear clustering by phage genus, with phage families also forming distinguishable groups **(Fig. 3A)**. Using the RB-TnSeq matrix of the high-scoring fitness genes, we generated Bray-Curtis dissimilarity matrices and visualized the data using Principal Coordinates Analysis (PCoA) and Non-metric Multidimensional Scaling (NMDS) ordination methods (**Fig. 3BC**). Pairwise PERMANOVA analyses confirmed that phages clustered significantly according to their taxonomy at the family and genus levels (**Tables S3–S5**) with one exception at the genus level that is likely attributable to the high genomic similarity between the two genera in question (48).

**Figure 3.**
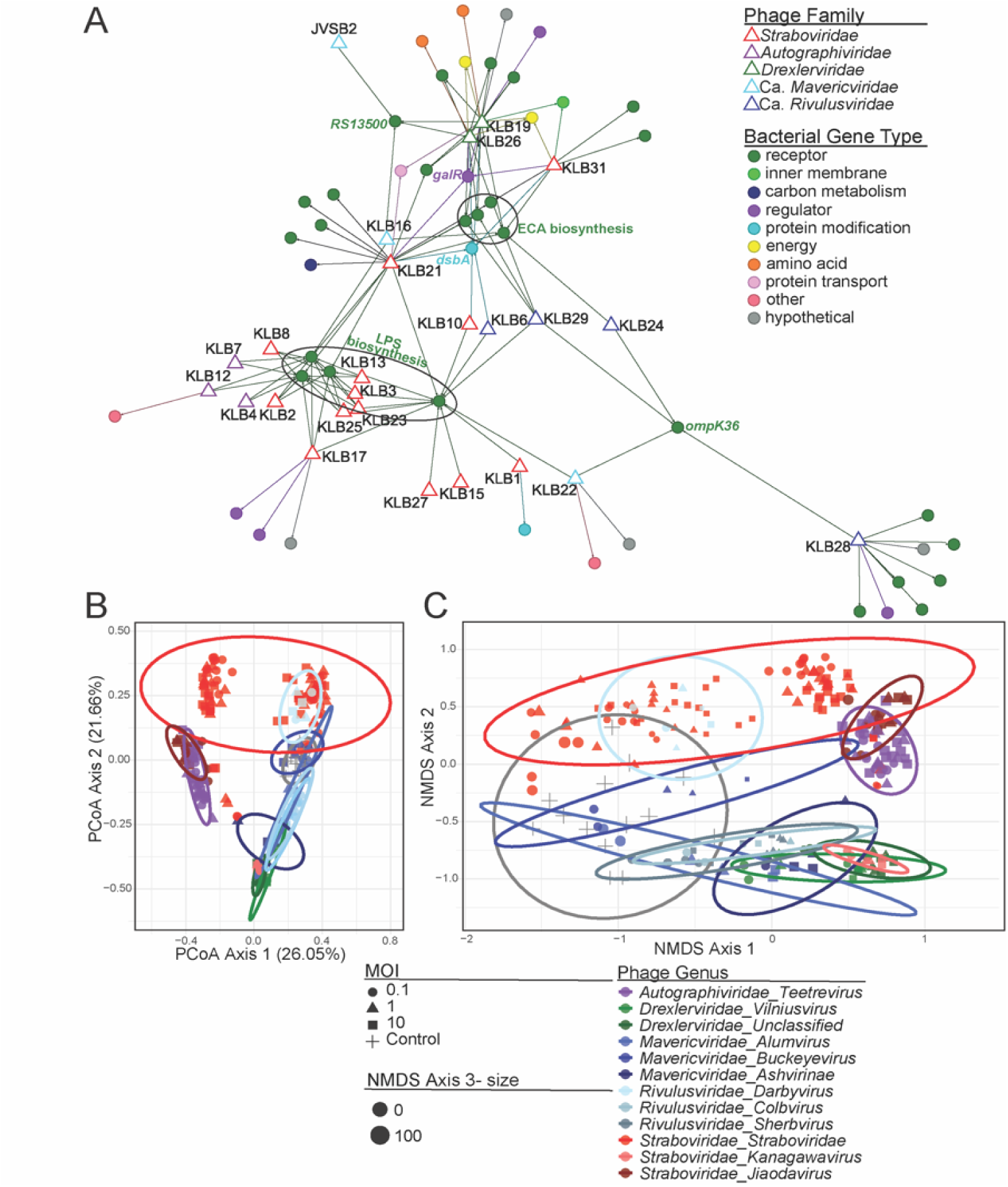
Patterns of phage infection. (A) Summary of bacterial gene requirement patterns between phages observed in our RB-TnSeq assays. Mixed-node network graph shows connections between phages (open triangles) and bacterial genes (solid circles). Gene nodes are colored by function (see legend), and bacterial genes that provide broad cross-resistance (i.e, to four or more phages) are labeled. (B) Principal Coordinates Analysis (PCoA) and (C) Non-metric Multidimensional Scaling (NMDS) plots based on Bray-Curtis dissimilarity of fitness profiles across samples. Each point represents a sample, colored by phage genus and shaped according to the multiplicity of infection (MOI). Ellipses indicate 95% confidence intervals for each genus. In the NMDS plot, point size corresponds to variation along the third NMDS axis.

At the family level, all tested families exhibited significant clustering (p_adjusted_ < 0.05), indicating that phage families differ consistently in their host gene usage patterns. Where multiple genera were available for testing, genus-level comparisons consistently showed stronger statistical support and the highest R^2^ values than the corresponding family-level comparisons (**Table S3-S5**), demonstrating that genus is a stronger predictor of host gene requirements for phage infection than family. Overall, our results suggest that phage genus membership carries quantifiable predictive information about infection strategy, and phylogenetic classification at genus resolution can inform host dependency modeling.

The divergence between genus-level and family-level predictive power was not uniform across all groups. Within the *Straboviridae*, *Slopekvirus* phages share the majority of their genome content divided into two distinct receptor dependency profiles, a pattern not resolvable at the family level. This suggests that within well-established families, genus-level resolution is necessary to capture meaningful variation in infection strategy.

Whether this relationship between taxonomic resolution and predictive power generalizes across broader *Klebsiella* phage diversity, and whether the novel families Ca. *Mavericviridae* and Ca. *Rivulsuviridae* show distinct scaling behavior given their greater phylogenetic distance from the established families, remains to be determined as collections of sequenced isolates in these groups expand. For now, the data support genus as the important unit of functional prediction within the families represented here, while acknowledging that the optimal taxonomic resolution for predicting infection strategy may vary across the broader phage diversity infecting this host.

### Receptor gene dependencies and intra-genus diversification

The majority of the 25 putative receptor hits encode genes involved in the biosynthesis of conserved surface structures, consistent with prior observations that phages preferentially target abundant, evolutionarily conserved surface features (56). Among the surface-associated hits, the largest groups were enterobacterial common antigen biosynthesis (ECA) (n = 7), cellulose biosynthesis (n = 5), LPS biosynthesis (n = 4), and genes involved in both LPS and ECA biosynthesis (n = 4), alongside the individual loci *tolC*, *ompK36*, *fadL*, *pqqU* and two poorly characterized genes, RS16715 and RS24775 (**Fig. 2**). RS16715 encodes a protein containing a DUF535 domain with homology to VirK, a family associated with outer membrane integrity and virulence in gram-negative bacteria, while RS24775 encodes a predicted inner membrane protein with homology to YqjF, a poorly characterized protein of unknown function. The roles of these two genes in phage infection have not been previously described and represent novel candidates for future characterization. Four of these groups, ECA, LPS, *ompK36*, and *pqqU* conferred cross-resistance to four or more phages across at least two phage families (**Fig. 2**). This finding is consistent with previous studies showing that bacterial genes involved in phage adsorption such as those involved in LPS biosynthesis often provide cross-resistance across multiple phage families, and with some phages binding to more than one receptor (39,40).

Enterobacterial common antigen (ECA) biosynthesis genes represented the largest single receptor-associated group in the screen, with seven genes required across multiple phages spanning at least two families. ECA is a conserved glycolipid present on the outer membrane of virtually all Enterobacteriaceae (57), and ECA biosynthesis genes have been identified as phage receptor determinants in genome-wide screens in *E. coli* and *Salmonella* (43), though its role in *Klebsiella* phage infection had not been previously described. The ECA-dependent phages in this dataset span more than one family, suggesting that exploitation of this surface structure has arisen independently across phage lineages, consistent with the general principle that abundant, conserved surface structures are convergently targeted by diverse phage groups (56). Several bacterial mutants in both ECA and LPS biosynthesis pathways showed elevated fitness scores for the same phages, raising the possibility that these phages can engage either surface structure and that receptor redundancy may buffer against resistance arising from disruption of a single biosynthesis pathway. Whether these hits reflect direct ECA-dependent binding or dependence on a related glycan produced by the same biosynthetic pathway, as recently described in *E. coli* (43), remains to be determined in *Klebsiella*.

Five genes involved in cellulose biosynthesis also showed high fitness score, all associated with a subset of phages from the same family. Cellulose is an extracellular matrix component produced by many soil-dwelling bacteria, including *Klebsiella* species, where it contributes to biofilm formation and surface attachment (58). Its presence on the bacterial surface makes it a structurally accessible candidate receptor, and cellulose has been identified as a phage receptor in other gram-negative bacteria (59,60). Given that *Klebsiella* sp. M5al is a soil isolate in which cellulose production is likely ecologically relevant; these hits may reflect receptor interactions that are particularly important in environmental rather than clinical contexts, where cellulose-producing biofilm communities predominate.

To examine how receptor diversity arises within a single genus, we focused on the *Slopekvirus* genus, the largest group in our collection with ten phages that share 199 of an average of 277 genes per genome. Despite this high genomic similarity, the phages separated into two distinct receptor dependency profiles: all ten required *wabO*, but six additionally required *wabI*, *wabN*, and *wabH*, indicating differences in the specific LPS outer core regions recognized (**Fig. 4A,B**).

**Figure 4.**
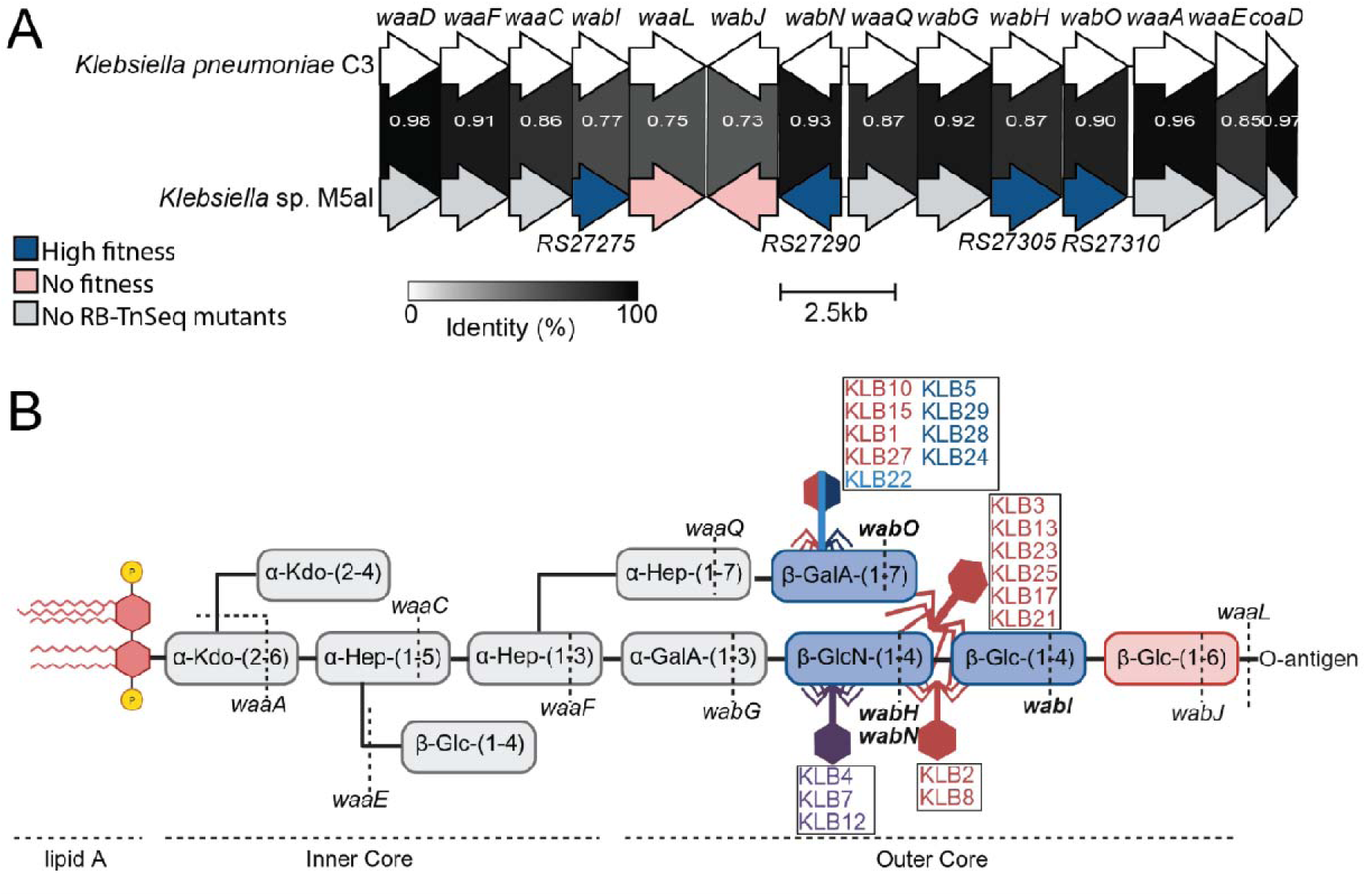
Impact of lipopolysaccharide (LPS) on phage infection. (A) Representation of the three operons involved in LPS biosynthesis. Bacterial genes are colored as follows: putative essential genes (colored gray), genes with high and significant fitness scores (fitness score > 4, blue), and genes that did not have significant fitness scores (pink). The gene names RS27275, RS27290, RS27305, and RS27310 were inferred by comparison to the previously characterized strain *K. pneumoniae* C3. Operons were visualized using Clinker. Links between genes denote percent amino acid identity between deduced proteins. B) Biochemical context showing the role of the LPS operon in the synthesis of the LPS core, and the RB-TnSeq-inferred interaction with phages. Phages are represented at the inferred binding region based on RB-TnSeq data, and color-coded by the phage family.

Comparison of receptor-binding protein sequences offered a structural explanation for this divergence. In *Straboviridae* phages, the long tail fiber tip is responsible for receptor binding (61), and amino acid comparisons of the distal tail fiber subunit across all ten *Slopekvirus* phages revealed high conservation in the N-terminal approximately 160 residues but substantial divergence in the C-terminal receptor-binding region (**Fig. 5A,B**). The two receptor-use profiles clustered by C-terminal sequence identity, and the receptor-binding domain of KLB25 showed greater similarity to the *wabI/wabN/wabH/wabO*-requiring group than to the *wabO*-only group, suggesting potential horizontal acquisition of this region consistent with its LPS binding profile. These data indicate that intra-genus divergence in receptor usage is driven by sequence variation in the distal tail fiber, and that subtle genetic variation among closely related phages can produce functionally distinct host interactions.

**Figure 5.**
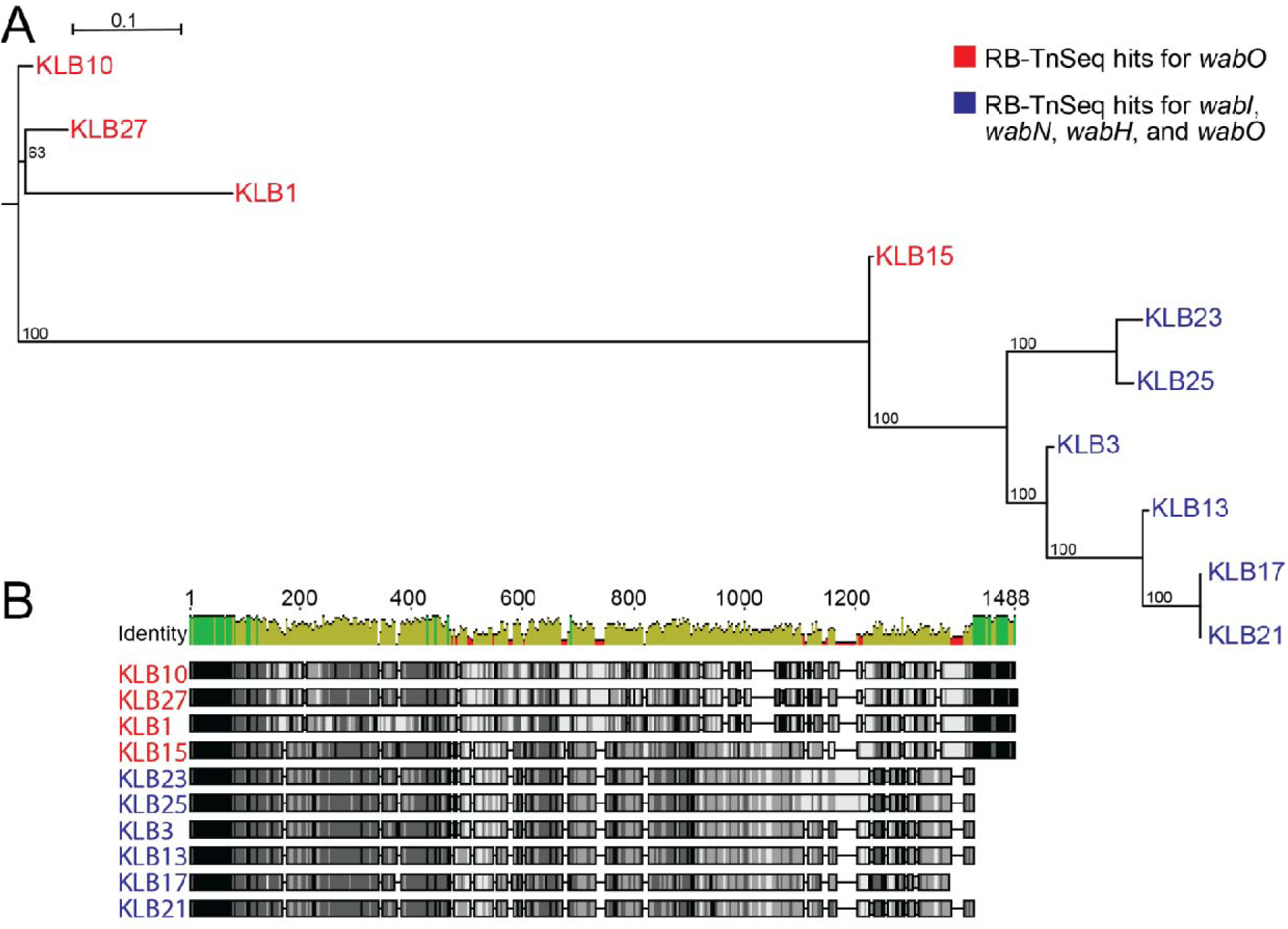
Genomic and structural differences in the long tail fiber of *Slopekvirus* phages correlate with LPS binding profiles. (A) Phylogenetic tree of the distal tail fiber subunit from *Slopekvirus* phages. Phages requiring only *wabO* for infection are labeled in red; phages requiring *wabI*, *wabN*, *wabH*, and *wabO* are labeled in blue. (B) Multiple sequence alignment of the distal long tail fiber subunit across all *Slopekvirus* phages, highlighting conserved and variable regions associated with receptor binding specificity.

Receptor usage and phage genus have emerged as key predictors of phage infectivity at the strain level (28,62–64). The present findings reinforce this view, further demonstrating that even phages sharing high genomic similarity within the same genus can rely on distinct receptor structures, with divergence in receptor-binding proteins as the primary driver. This observation aligns with experimental evolution studies showing that minor genetic changes can substantially alter host specificity, including single amino acid substitutions in receptor-binding proteins that redirect phage adsorption to different surface structures (20,22,44,65). Broad host-range screens consistently support this model across phage families (43,52,66–75).

### Non-receptor host factors uncovered in RB-TnSeq screens

Beyond surface receptor genes, we identified 17 bacterial genes involved in intracellular functions, the majority of which were phage-specific, associated with only one to three phages (**Fig. 2**). These genes generally produced lower fitness scores than the surface receptor hits, suggesting a more modest or indirect contribution to phage infection rather than an essential receptor role. These non-receptor dependencies span transcriptional regulation, inner membrane integrity, energy metabolism, protein folding, and amino acid transport. While none of these genes have been directly associated in the context of phage infection in *Klebsiella*, their known roles in cell envelope physiology and bacterial physiology provide a basis for hypothesizing how their disruption might influence phage infection.

The cytochrome bd oxidase subunits *cydA* and *cydB* were associated with phages KLB19, KLB26, and KLB31, all of which also produced fitness hits for *dsbA*. Given that cytochrome bd oxidase contributes to the reducing power required by DsbA (76,77), this co-occurrence suggests a potential functional link between these genes during infection. The transcriptional regulator SdiA was associated with phage KLB17; disruption of *sdiA* in *Klebsiella* leads to increased biofilm formation, impaired cell division, and filamentous morphology (78), any of which could alter phage adsorption. The inner membrane proteins NlpD and TolR, associated with phages KLB19, KLB21, and KLB31, are both implicated in cell envelope architecture. NlpD activates peptidoglycan amidases during cell division (78), and its disruption produces outer membrane defects that could affect phage binding or lysis. TolR is a component of the Tol-Pal system, which spans the inner membrane and periplasm and maintains outer membrane integrity; mutations in Tol-Pal genes in *E. coli* disrupt LPS structure (79), suggesting an indirect effect on phage receptor accessibility rather than a direct role in DNA translocation for the *Straboviridae* phages in this study. The core component of the twin-arginine translocation system *tatC* mutants showed fitness in presence of KLB19, KLB21, and KLB4. To assess whether the *tatC* association could reflect direct Tat-dependent export of a phage protein or screen hit, all phage-encoded proteins and high-fitness bacterial genes were systematically examined for canonical Tat signal motifs using SignalP-6.0. None were identified (**Table S6**), ruling out the simplest model and implicating a more indirect role for the Tat pathway, potentially through export of host proteins that maintain surface accessibility for the relevant phages.

### Experimental validation of RB-TnSeq hits

To confirm that RB-TnSeq fitness scores reflect genuine phage resistance phenotypes, we generated individual, sequence-confirmed knockout strains for eight genes spanning receptor biosynthesis, outer membrane transport, and regulatory functions. These include the LPS outer core biosynthesis genes *wabI* (RS27275), *wabN* (RS27290), and *wabH* (RS27305); the outer membrane porin *ompK36*; the outer membrane long-chain fatty acid transporter *fadL*; the pyrroloquinoline quinone biosynthesis gene *pqqU*; the predicted inner membrane protein RS24775; and the periplasmic disulfide bond catalyst *dsbA*. Complementation was performed for knockouts to exclude polar effects on neighboring genes.

### Validated surface receptor determinants

#### LPS outer core biosynthesis genes

The RB-TnSeq screen identified *wabI*, *wabN*, and *wabH* as conferring resistance to a subset of *Slopekvirus* phages (**Fig. 4AB**, **Fig. 6A)**. Disruption of each gene conferred complete resistance to these phages, with the exception of partial resistance observed in *Autographiviridae* phages, consistent with potential alternative receptor usage (**Fig. 6A**). Adsorption assays confirmed that phage binding was abolished in all three knockout backgrounds, directly supporting the role of these LPS outer core biosynthesis genes as functional phage receptor determinants (**Fig. 6B,C,D**). Complementation of each knockout restored wild-type infectivity, confirming that resistance was attributable to loss of the specific gene rather than polar effects on neighboring loci. These genes had not been characterized as phage receptors in any *Klebsiella* strain prior to this study. Repeated attempts to generate a clean deletion of *wabO* were unsuccessful; this hit is therefore retained as a high-confidence candidate for future validation.

**Figure 6.**
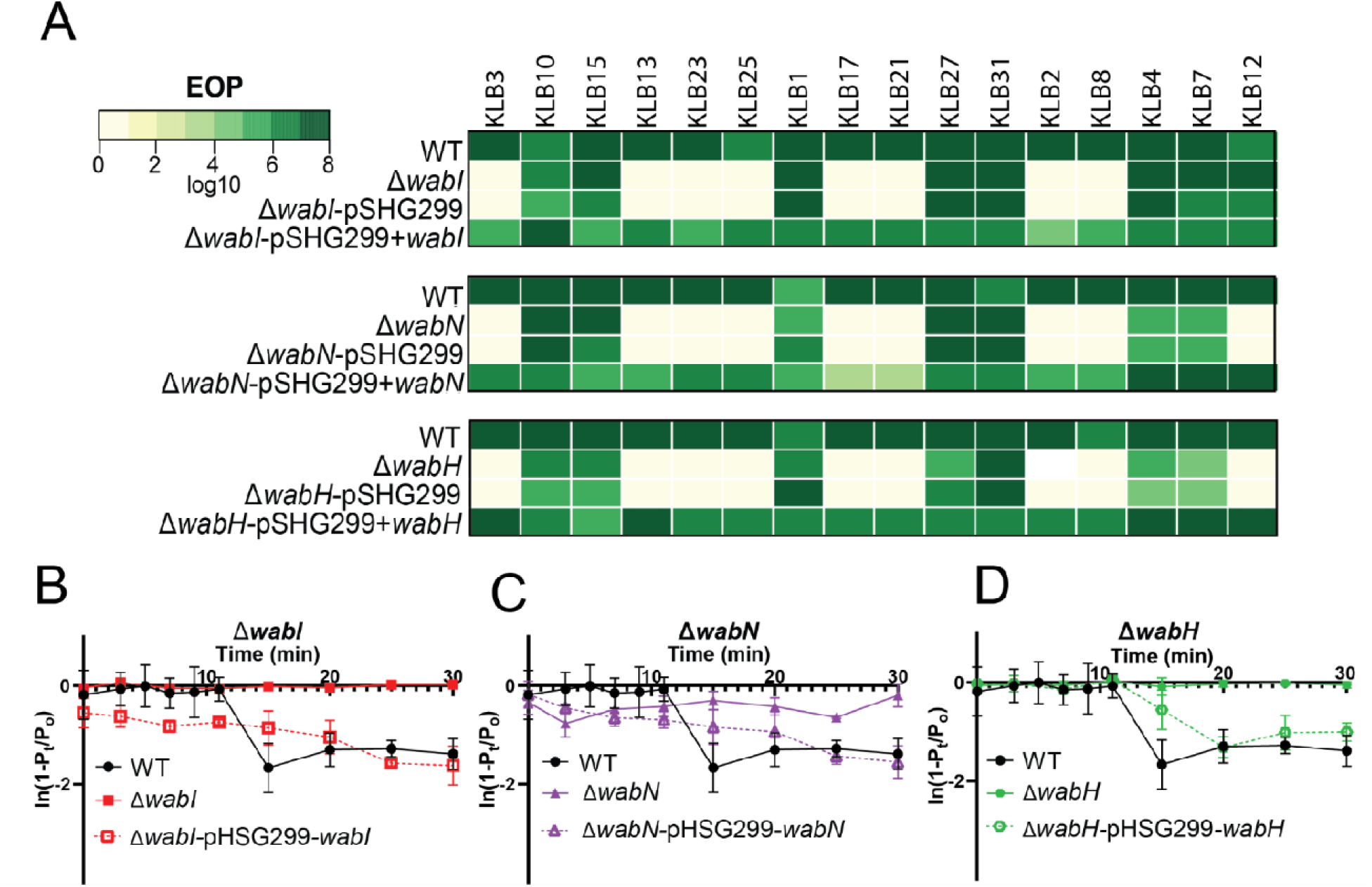
Impact of lipopolysaccharide (LPS) biosynthesis gene knockouts on phage infection. (A) Titers of the phages that required LPS to infect WT, ΔRS27275, ΔRS27290, and ΔRS27305, each of these mutants with the empty vector, and each mutant with a complemented gene. Black denotes phages not assayed as they did not require LPS based on the RB-TnSeq assay. Rate of adsorption of phage KLB2 for each of the three LPS mutants: (B) RS27275, (C) RS27290, and (D) RS27305. In adsorption assay panels (B-D), the solid black line represents the phage-only (no-bacteria) control, showing baseline phage titer in the absence of bacterial adsorption.

#### OmpK36

The screen identified *ompK36* as conferring cross-resistance across multiple phage families. A Δ*ompK36* knockout was generated and tested by efficiency of plating against KLB24, KLB28, and KLB29 phages with a corresponding high fitness score. Disruption of *ompK36* conferred strong resistance to all phages spanning two phage families, and complementation restored phage infectivity, confirming OmpK36 as a functional cross-family receptor in *Klebsiella* sp. M5al (**Fig. 7A**). OmpK36 was previously identified as a receptor in a capsulated *K. pneumoniae* strain, where it functioned as a secondary receptor accessed after capsule depolymerization (33). Its validation here as a primary receptor in an acapsular host directly demonstrates that the receptor landscape exposed in the absence of a capsule is both distinct from and complementary to that characterized in capsulated strains.

**Figure 7.**
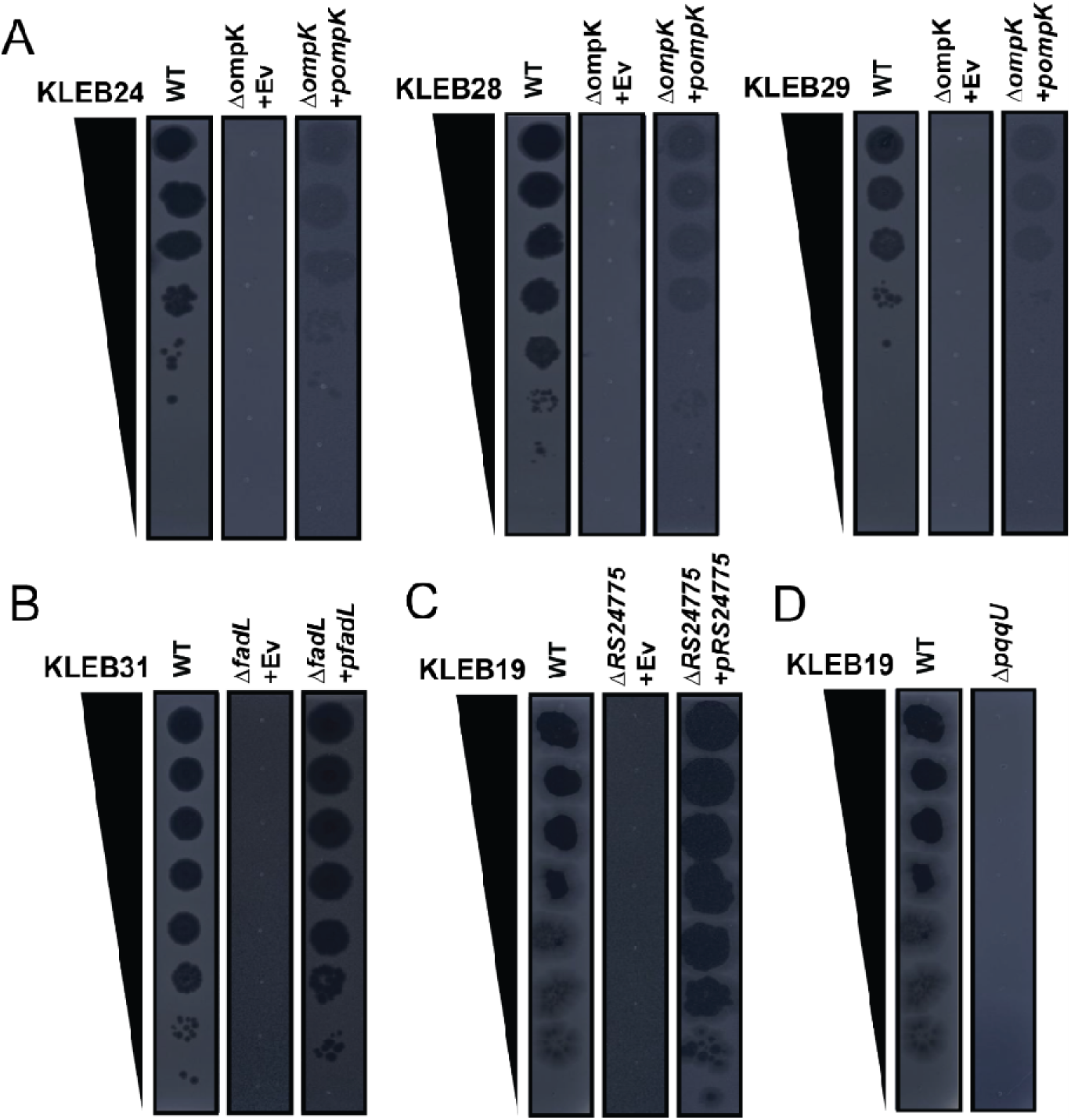
Experimental validations of top scoring RB-TnSeq hits. Efficiency of plating for phages with RB-TnSeq hits in (A) Δ*ompK36*, (B) Δ*fadL*, (C) Δ*RS24775 (D)* Δ*pqqU.* WT is wild type host strain, Ev is an empty vector, p is for complementation plasmid.

#### FadL

The screen identified *fadL*, encoding an outer membrane long-chain fatty acid transporter, as a hit for phage KLB31, a member of the *Straboviridae*. A Δ*fadL* knockout was generated and infectivity assessed by efficiency of plating. Disruption of *fadL* conferred resistance to KLB31, and complementation restored wild-type infectivity, identifying FadL as a functional receptor determinant for this phage (**Fig. 7B**). FadL has been previously identified as a phage receptor in *E. coli* (39,43), and this work represents the first characterization of *fadL* as a receptor in *Klebsiella*.

#### PqqU

The screen identified *pqqU*, involved in pyrroloquinoline quinone biosynthesis, as a hit for phages KLB19 and KLB26, both members of the *Straboviridae*. A Δ*pqqU* knockout was generated and tested by the efficiency of plating against KLB19. Disruption of *pqqU* conferred resistance to KLB19 (**Fig. 7C)**. The role of PqqU in phage infection had not been previously described in *Klebsiella*, but has been found in other phage-host systems (43,80). Its identification here raises the possibility that PQQ-dependent surface modifications influence receptor accessibility for a subset of *Straboviridae* phages. Notably, while *pqqU* was required for both KLB19 and KLB26, *RS24775* was identified as an additional hit specific to KLB19 (see below) but not KLB26, suggesting that KLB19 may depend on a broader set of host factors for productive infection than KLB26, despite both phages sharing the *pqqU* requirement. Whether RS24775 and PqqU act in the same or independent pathways during KLB19 infection represents a question for future investigation.

### Validated non-receptor host factors

#### RS24775

The screen identified RS24775, encoding a predicted inner membrane protein with homology to YqjF, as a hit for phage KLB19, a *Straboviridae* phage. A Δ*RS24775* knockout was generated and infectivity assessed by efficiency of plating. Disruption conferred resistance to KLB19, and complementation restored wild-type infectivity (**Fig. 7D)** indicating RS24775 role in KLB19 phage infection. The function of YqjF-family proteins is not well characterized in any organism, and this represents the first reported association of this protein family with phage susceptibility. Its inner membrane localization suggests a possible role in post-adsorption infection steps rather than direct receptor function, though this awaits experimental investigation.

#### DsbA

The screen identified *dsbA* as a hit for seven phages across the *Straboviridae* and *Drexlerviridae* families. The fitness signal for *dsbA,* while reproducible across multiple independent transposon insertions, was lower in magnitude than that observed for the receptor genes described above, predicting partial rather than complete resistance. To test this prediction, plating efficiency and liquid growth inhibition were compared in wild-type versus Δ*dsbA* cells. The Δ*dsbA* mutant did not confer complete resistance, with efficiency of plating measurements revealing modest reductions for a subset of phages (**Fig. S1**), and liquid growth assays showing a significant decrease in infectivity for phage KLB1 in the Δ*dsbA* background while other phages showed no significant difference (**Fig. S1**). DsbA catalyses disulfide bond formation in periplasmic and outer membrane proteins, and its disruption is known to impair the folding and surface presentation of multiple outer membrane proteins in gram-negative bacteria. In *Klebsiella* sp. M5al, where LPS outer core and outer membrane porins serve as primary receptor structures, DsbA-dependent folding of one or more surface components could plausibly influence their accessibility or structural integrity as phage receptor targets, accounting for the partial resistance phenotype observed across multiple phage families.

The co-occurrence of *cydA* and *cydB* fitness signals across the same phages is consistent with this interpretation, as cytochrome bd oxidase contributes to the reducing power that supports DsbA activity (76). The attenuated Δ*dsbA* phenotype may additionally reflect functional redundancy with the disulfide bond isomerase DsbC, which cooperates with DsbA in maintaining periplasmic protein folding and could partially compensate for DsbA loss in certain contexts (76). A prior transposon screen in *Acinetobacter* also identified *dsbA* as a phage infection determinant, where its disruption truncated the capsular polysaccharide and paradoxically increased phage adsorption (81). Additionally in phage lambda assays, *dsbA* disruption was found to have a moderate delay in lysis timing (82). In *Klebsiella* sp. M5al, which is naturally acapsular, this capsule-dependent mechanism cannot apply, and the opposing direction of the phenotype, namely partial resistance rather than increased susceptibility, suggests that DsbA influences phage infection through a distinct, capsule-independent pathway in this host. Whether a common surface-remodelling regulatory mechanism underlies DsbA effects across both systems represents an open question for future investigation.

Collectively, the 42 genes identified in this work reveal the breadth of host functions co-opted by diverse *Klebsiella* phages, from surface receptor biosynthesis to intracellular metabolism, and provide a resource for directing studies of phage-host interaction mechanisms (**Fig. 2**, **Fig. 8, Table S7).**

**Figure 8.**
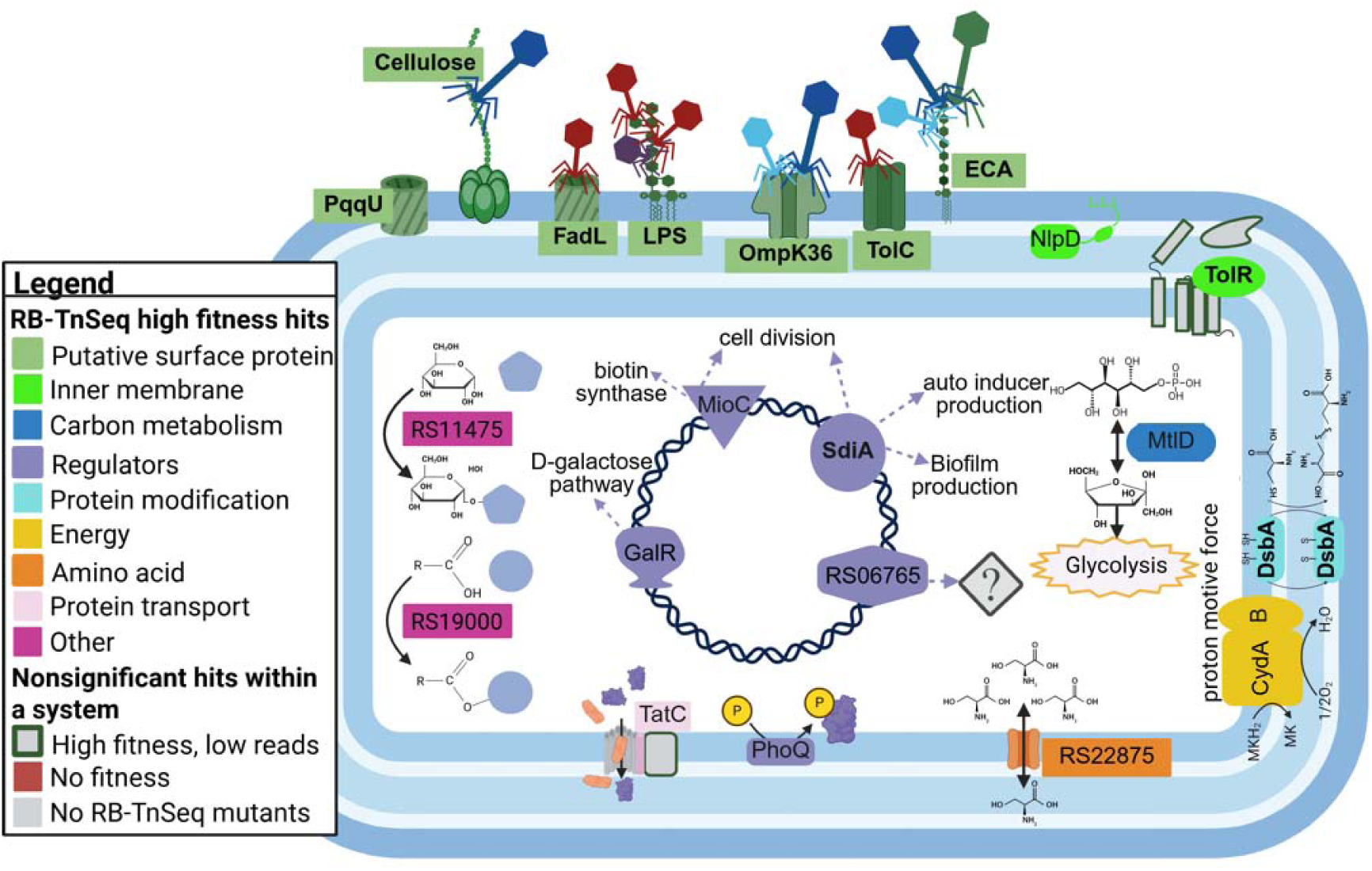
Overview of phage-bacteria interactions during phage infection. A total of 42 bacterial genes were identified in the RB-TnSeq screen. Bacterial genes are color-coded by their predicted function: surface receptor (green), inner membrane protein (light green), carbon metabolism (blue), regulators (purple), protein folding (light blue), energy production (yellow), protein transport (light pink), and other proteins (dark pink). For bacterial genes that are part of a complex, the complex is color-coded based on RB-TnSeq results. Green border: high fitness but read count less than 100; gray: no RB-TnSeq mutants available.

## Conclusion

Our study delivers the first genome-wide, cross-family map of phage infection dependencies in *Klebsiella*, built deliberately on an acapsular host that reveals the receptor landscape invisible to prior capsule-centric studies. Systematic characterization of phage receptors in *Klebsiella* has been largely unexplored with decades of individual phage studies in capsulated strains that collectively produced knowledge of a handful of receptors for a small number of phages (28,66–68,83). By examining phage infection in a non-capsulated host, we uncovered over 42 high scoring broader sets of host determinants that influence phage infection and are likely obscured in capsule-centric studies. These results expand our understanding of phage-host interactions and highlight overlooked genetic factors that may govern resistance and host range.

To move from descriptive studies to predictive frameworks, we must identify conserved infection strategies across taxonomic boundaries. Our findings show that bacterial gene requirements for infection significantly cluster by phage genus and family, suggesting that infection strategies are often taxonomically structured. However, even within these groups, we observed divergences in host gene dependencies, particularly in receptor usage and intracellular processes. This variation underscores the value of pairing phage genomes with systematic host genetic screens like RB-TnSeq. This type of classification provides a functional layer of prediction that goes beyond sequence similarity and can help us understand the mechanisms of host-range.

In summary, this work advances the understanding of *Klebsiella* phage-host interactions by revealing the receptor architecture beneath the capsule, and provide a community resource for linking phage genomics to functional host interactions. These findings support the rational design of phage cocktails that account for capsule-loss escape variants, inform host-range prediction, and contribute to the broader goal of moving from descriptive phage genomics to mechanistically grounded, predictive models of phage infectivity.

## Methods

### Bacterial strains and growth conditions

Bacterial strains, phages, and their sources are listed in (**Table S8**). *Klebsiella* sp. M5al was grown in LB broth at 30°C, with shaking at 150 rpm. All *E. coli* strains used for plasmid maintenance and conjugation were grown in LB at 37°C and shaking at 150 rpm. All bacterial strains were stored long-term in –80°C with 15% glycerol.

### Bacteriophages and propagation

The *Klebsiella* sp. M5al phages were isolated, annotated, and 24 of the 25 phages described by our group previously (48). Phages were amplified using standard protocols (107). Briefly, *Klebsiella* sp. M5al was grown to mid-exponential (OD of 0.3) and five phage plaques were added to the culture. The infected culture was then incubated overnight, and the lysates were centrifuged and filtered using a 0.22 µm filter. Phage titer was determined by diluting lysates in SM buffer (100mM NaCl, 8mM MgSO_4_, 50mM Tris, pH 7.5) and mixing 100 µL of diluted phages with 3 mL of 0.7% LB agar supplemented with 10 mM CaCl and 10 mM MgSO, and 200 µL of overnight culture. Plates were incubated at 30 °C overnight.

### Competitive growth experiments of the RB-TnSeq library

*Klebsiella* sp. M5al RB-TnSeq library was previously created by collaborators at the Lawrence Berkeley National Laboratory (33). Briefly, the library was generated using the Tn5 transposase and consists of 134,723 unique barcodes. There were, on average, six knockouts per nonessential gene. A single aliquot (1 mL) of *Klebsiella* sp. M5al RB-TnSeq library stored at –80 °C was thawed and transferred to LB supplemented with 50 µg/mL of kanamycin. The culture was grown to mid-exponential at 30 °C with shaking at 150 rpm until the OD reached between 0.2-0.5, measured with an Epoch 2 plate reader. From this culture, three 1-mL aliquots were sampled and spun at 21,130 x g to pellet. These bacterial pellets were used as the reference for BarSeq (termed T0). The library was then diluted to an OD of 0.04 and dilute in LB medium prepared at 2× concentration, containing 20 g tryptone, 10 g yeast extract, and 10 g NaCl per liter. In a 48-well plate, 350 µL of the diluted RB-TnSeq mixture was then mixed in a 1:1 ratio with 350 µL of phages diluted in SM buffer at three MOIs (0.1, 1, and 10). The plate was covered with a Breathe-Easy film, and the 48-well plate was then placed in an Epoch 2 plate reader at 30 °C with double orbital shaking at 150 rpm and OD 600 nm readings every 15 minutes for 4.5 hours. At the end of the experiment, each well contents were transferred to a 1.5-mL microcentrifuge tube and spun at 21,130 x g to pellet the bacterial cells. The supernatant was removed, and the bacterial pellets were placed at – 80 °C until DNA was isolated for BarSeq sequencing.

For each phage, there were at least two biological replicates for each time point, with additional replicates sometimes performed for clarity (**Table S9**). In total, there were 213 phage-infected RB-TnSeq assays. Across these 213 assays, we identified 669 significant fitness hits (**Table S10**) that dispersed across 63 bacterial genes. These hits were further curated based on read abundance (>100 reads for three strains) to reduce the chances of false positives (n = 16) and checked for potential polar effects (n = 5), resulting in 42 bacterial genes with significant fitness changes (**Fig. 2, Table S11**).

### BarSeq of RB-TnSeq pooled fitness assay samples

We isolated genomic DNA from RB-TnSeq library samples using the DNeasy Blood and Tissue kit (Qiagen). We performed 98°C BarSeq PCR protocol as described previously (38,39) with the following modifications. Barcodes were PCR amplified using “BarSeq_V4” primers, as described previously (34). These included a 10-bp index sequence for P7 primers and an internal 8-bp index (to identify Illumina index hopping), which allowed multiplexing up to 8 plates of 96-well BarSeq PCR samples (total 768 genome-wide assay samples). All experiments done on the same day and sequenced on the same lane were considered as a ‘set’. Equal volumes (5 µl) of the individual BarSeq PCRs were pooled, and 50 µl of the pooled PCR product were purified with the DNA Clean and Concentrator kit (Zymo Research). The final BarSeq library was eluted in 40 µl of sterile water.

### Data processing and analysis of BarSeq reads

BarSeq reads were converted to counts per barcode using the MultiCodes.pl script in the feba code base (https://bitbucket.org/berkeleylab/feba/), with the –minQuality 0 option and –bs4 (BarSeq 4) for BarSeq primers. RB-TnSeq fitness data was analyzed as previously described (38,39). Briefly, the fitness value of each strain (an individual transposon mutant) was the normalized log2(strain barcode abundance at end of experiment/strain barcode abundance at the start of the experiment). The fitness value of each gene was the weighted average of the fitness of its strains. Because RB-TnSeq transposons were redundant, with on average six barcodes interrupting each nonessential gene and a minimum of at least three barcodes for each gene in our screen, the fitness score for each gene was determined by averaging the fitness levels of these strains. A positive fitness score indicated that the knocked-out host gene increased the relative fitness of the strain in the presence of phages, suggesting that the gene product was necessary for successful phage infection. Conversely, a negative fitness score suggested that the knocked-out strain was more susceptible to phage infection. We applied filters on experiments such that the mean reads per strain were >= 10. In a typical experiment without stringent positive selection, a gene fitness score (fit) > 1 and an associated t-like statistic >4 sufficed to give a low rate of false positives. Due to the stringent positive selection in phage assays, we required that fit ≥ 4, t ≥ 5, standard error = fit/t ≤ 2, and fit ≥ maxFit – 8, where maxFit is the highest fitness value of any gene in an experiment. The limit of maxFit – 8 was chosen based on our earlier work (39).

### Annotation of bacterial genes identified in RB-TnSeq

To understand the functional significance of the 42 identified bacterial genes required for phage infection, we first classified them into nine functional categories-surface structures, inner membrane proteins, transcriptional regulators, carbon metabolism, energy cycling, amino acid biosynthesis and transport, protein transport, protein modification, other genes such as glycosyltransferase and acyltransferases that do not have specified pathways, as well as genes consisting of hypothetical proteins with no known function (**Fig. 2**). The first eight functional categories were relatively straightforward to interpret. In contrast, the latter “other” category and hypothetical proteins were more difficult to analyze.

To refine our understanding of these latter two categories, the thirteen genes were more deeply assessed via homology-based searches, domain analyses, protein structure predictions, operon predictions, and constructed protein-protein interaction networks (**Fig. S2; Tables S12-S13**) (84–88). Together, these analyses provided complementary lines of evidence to generate putative functional annotations for otherwise uncharacterized proteins.

Five of these bacterial genes did not have enough evidence to associate with a specific gene or functional pathway. *RS27275*, *RS27305*, and *RS27310* were identified as family 1 glycosyltransferases, which mediate the transfer of sugar moieties to form glycosidic bonds. This glycosyltransferase family involves several pathways, including exopolysaccharide biosynthesis, colanic acid production, and lipopolysaccharide (LPS) biosynthesis. Notably, *RS27275, RS27290, RS27305,* and *RS27310* were clustered within eight genes of one another and were organized into three consecutive operons within the bacterial genome, all of which were associated with LPS biosynthesis. Multiple Sequence Alignment (MSA) confirmed that these genes were >80% similar to known homologs (89,90), and they were renamed respectively RS27275-*wabI*, *RS27290*-*wabN*, *RS27305*-*wabH*, and *RS27310*-*wabO*. Additionally, RS18965, RS18970, RS18985, and RS18990 were predicted to play a role in O-antigen and enterobacterial common antigen (ECA) biosynthesis and were placed in the surface structure category.

To see if some of the proteins had TAT domains, indicating these proteins are exported via the Tat system, we used SignalP-6.0 to identify proteins with the TAT domains.

### Generation of individual *Klebsiella* sp. M5al knockouts

All individual Klebsiella sp. M5al single gene deletions were created using one of two approaches depending on the gene target.

The majority of gene deletions were created using the VcDART, a CRISPR-associated transposase DART system, to make targeted knockouts (91).The gRNAs are 32 nucleotides long and recognize targets with a 5′-CC-3′ type I-F protospacer adjacent motif (PAM). For each deletion, gRNAs were designed targeting the first half of the gene of interest to ensure functional knockouts after transposon insertion (**Table S14**) with additional nucleotides added at the 5’– and 3’-ends to generate sticky ends for insertion into plasmid pBFC0619.

Oligos were inserted into plasmid pBFC0619 using a previously reported method involving the BsaI-HFv2 Golden Gate assembly kit (NEB). Briefly, 5 µL of the Golden Gate assembly reaction was mixed with 50 µL of competent *E. coli* and subjected to heat shock at 42 °C for 60 seconds, followed by immediate placement on ice for 5 minutes. Then, 1 mL of SOC broth (Super Optimal broth with Catabolite repression) was added, and cells were incubated at 37 °C with shaking at 150 rpm for 60 minutes before plating on LB agar supplemented with 10 µg/mL gentamicin. SOC broth consisted of 2% (w/v) tryptone, 0.5% (w/v) yeast extract, 10 mM NaCl, 2.5 mM KCl, 10 mM MgCl, 10 mM MgSO, and 20 mM glucose. gRNA insertion was confirmed via PCR and Sanger sequencing.

For gRNA plasmids that only had high efficiency of transformation in pir+, the plasmid was isolated using Qiagen plasmid mini extraction kit. A 10 mL culture of *E. coli* pir+ transformant was grown in LB supplemented with 10 µg/mL of gentamicin at 37 °C with shaking at 150 rpm. The 10 mL of culture was then pelleted, and the plasmid was isolated using the Qiagen plasmid mini extraction protocol. Plasmid was then electroporated into electrocompetent *E. coli* WM3064 (25 μF; 200 Ω, 2.5 kV), incubated for 2 hours at 37 °C with shaking at 150 rpm, and plated on 10 µg/mL gentamicin (92).

To generate the targeted knockout, the pBFC0619 plasmid with the inserted gRNA was transferred to *Klebsiella* via conjugation with *E. coli* WM3064 using previously established protocols (91). The donor and acceptor strains were mixed in 1:1 ratios on top of a 0.22 µm filter on an LB plate supplemented with DAP (0.06 mg/mL) and IPTG (0.1 M). IPTG induced the CRISPR-associated transposase to insert the transposon into the gene of interest. The plates were incubated at 30 °C for 6-8 hours. The filter was resuspended in 2 mL of LB and plated on LB supplemented with 25 µg/mL of gentamicin. Transposon insertion into the gene of interest was confirmed with Sanger sequencing.

For a subset of deletions (*pqqU*, *fadL*, *RS24775*), targeted knockouts were generated using the lambda Red recombineering system carried on plasmid pSim5 (93). Electrocompetent *Klebsiella* sp. M5al cells were prepared as follows (94). Cells were grown overnight to saturation, diluted 1:100 into fresh LB, and grown to an OD600 of approximately 0.1. The lambda Red system was induced by shifting cultures to 42°C and growth was continued until an OD600 of approximately 0.5 was reached. Cultures were immediately placed on ice for 15 to 30 minutes, washed three times with ice-cold 10% glycerol, and concentrated 100-fold in ice-cold 10% glycerol. A deletion cassette consisting of a kanamycin resistance marker flanked by sequences homologous to the regions immediately upstream and downstream of the target gene was amplified by PCR. Fifty microlitres of electrocompetent cells were electroporated with 50 ng of the PCR product using 0.1-cm cuvettes at 1.8 kV, recovered in 1 mL of LB for 2 hours at 30°C, and plated on LB agar with kanamycin selection at 30°C for 24 hours. Successful integration of the kanamycin cassette at the target locus was confirmed by whole-genome sequencing. All *Klebsiella* sp. M5al mutants were stored at –80°C in LB supplemented with 15% glycerol.

### Complementation of *Klebsiella* sp. M5al knockouts

Complementation experiments were performed using two approaches depending on the gene target.

For the majority of complementation constructs, the target bacterial gene was PCR-amplified using primers incorporating SbfI and SphI restriction sites (**Table S15**). The amplified DNA was gel-purified and digested alongside the plasmid vector pSHG299 using *SbfI* and *SphI* restriction enzymes (New England Biolabs). For ligation, 66 ng of the digested PCR product were combined with 33 ng of linearized pSHG299 (2:1 molar ratio), T4 DNA ligase, and the supplied buffer, and incubated under standard ligation conditions. The resulting ligation mixture was transformed into *E. coli* pir+ via heat shock. Transformants were screened by colony PCR to verify the presence of the insert. Plasmid DNA was isolated from positive clones using the QIAprep Spin Miniprep Kit (Qiagen) and electroporated into electrocompetent *Klebsiella* sp. M5al cells. Following electroporation, cells were recovered for 2 hours at 37 °C in 1 mL of SOC broth, then plated on LB agar containing 50 µg/mL of kanamycin. Successful plasmid integration into the genome was confirmed by colony PCR and Sanger sequencing.

For complementation of *RS24775* and *fadL*, a modified vector was constructed by replacing the kanamycin resistance marker in pSHG299 with a chloramphenicol resistance marker to enable selection in the (pSim5 enabled) kanamycin-marked deletion backgrounds. The target genes were PCR-amplified with primers incorporating overlapping sequences compatible with the modified vector. Amplified inserts were cloned into the linearized chloramphenicol-marked vector using the NEBioLabs Gibson Assembly mix according to the manufacturer’s instructions. The assembled constructs were electroporated into the corresponding deletion strains using the electrocompetent cell preparation protocol described above. Following electroporation, cells were recovered for 2 hours at 30°C in 1 mL of LB broth and plated on LB agar containing 300 µg/mL chloramphenicol. Successful construct integration was confirmed by colony PCR and Sanger sequencing. All complementation strains were stored at –80°C in 15% glycerol until use.

### Phage infection efficiency of *Klebsiella* sp. M5al knockouts

Phage resistance was assayed through efficiency of plating (EOP) experiments. Bacteria were grown overnight at 30 °C with shaking at 150 rpm and with 25 µg/mL of gentamicin. For *RS24775* and *fadL* deletion strains carrying empty or complementation constructs, overnight cultures were supplemented with both 50 µg/mL kanamycin and 300 µg/mL chloramphenicol to maintain both the deletion and complementation constructs simultaneously. An aliquot of 300 μL of the overnight culture was mixed with 5 mL of top agar (0.7% agar) and solidified at room temperature. Phages were diluted to a titer of 10^8^ PFU/mL and then ten-fold serially diluted in SM buffer. An aliquot of 5 µL of each phage dilution was then spotted onto the solidified bacterial lawn. The plates were incubated overnight at 30 °C. EOP was calculated as the ratio of the average effective phage titer on the *Klebsiella* sp. M5al knockout compared to the wild-type (WT) *Klebsiella* sp. M5al host. All plaquing experiments were performed with at least three biological replicates and with at least three overnight cultures for each bacterial strain.

### Phage mediated growth inhibition assays

Phage-mediated killing of *Klebsiella* sp. M5al was quantified using a plate reader–based growth inhibition assay. Bacterial cultures were grown to mid-exponential phase (OD 600nm = 0.3–0.6) and diluted in 2× LB to an OD of 0.1. MgSO_₄_ and CaCl_₂_ were supplemented to a final concentration of 20 mM each. Phages were diluted in SM buffer to a final titer of 4.1 × 10 PFU/mL, corresponding to a multiplicity of infection (MOI) of 6 prior to mixing with bacteria (resulting in a final MOI of 3 after combining equal volumes of phage and host).

Phage suspensions (350 µL) were aliquoted into a 48-well flat-bottom microtiter plate according to a predefined layout, followed by the addition of 350 µL of the diluted *Klebsiella* suspension, for a final volume of 700 µL per well. Plates were sealed with Breathe-Easy film and incubated in an Epoch 2 microplate reader. OD was measured kinetically every 5 minutes over 12 hours, with shaking at 205 cpm using a double-orbital setting between reads.

Growth inhibition was quantified by calculating the area under the curve (AUC) using GraphPad Prism. To account for baseline growth differences, AUC values from noninfected controls were subtracted from those of the corresponding phage-infected cultures. Statistical significance comparing phage infection of WT bacteria to Δ*dsbA* was done using a two-tailed *t*-test in GraphPad Prism. P values are denoted as follows: *P* < 0.05 (\**), P < 0.01 (**), and P < 0.001 **(\*\*\****).

### Examining the patterns of bacterial resistance

RB-TnSeq network graphs were constructed using Gephi. The python scripts to generate the Gephi graphs were previously described (36). In all graphs, edges were calculated based on datasets found in **Table S16**. Each mixed node graph in **Fig. 3** was drawn with weight one between a phage node (fixed) and a gene node if that gene conferred resistance.

To assess patterns in bacterial gene dependencies across phages, we performed Principal Coordinates Analysis (PCoA) and Non-metric Multidimensional Scaling (NMDS) using Bray-Curtis dissimilarity matrices. All analyses were conducted in R (version 4.3.0). RB-TnSeq gene fitness data were imported and transformed into matrix format. Since for viral RB-TnSeq we only look at the positive RB-TnSeq values as the negative fitness values were indicative of phage infection, we set negative values to zero to avoid distortion in similarity calculations.

For both the PCoA and NMDS, we calculated Bray-Curtis dissimilarity matrices using the vegdist function (117). PCoA was performed using cmdscale, and the percentage of variation explained by each axis was calculated from the eigenvalues. For NMDS, we used the metaMDS function (95) with three dimensions (k = 3) and a maximum of 100 iterations (trymax = 100). The NMDS stress value was used to assess model fit, with values <0.2 considered acceptable (96). The third NMDS dimension was scaled by a factor of 100 to be visualized as point size in ggplot2.

PCoA and NMDS coordinates were merged with sample metadata and visualized using ggplot2 (97), with colors corresponding to phage family, genus, or species, and ellipses representing 95% confidence intervals. To test for significant differences along PCoA axes, we performed Kruskal-Wallis tests followed by Dunn’s multiple comparisons with Bonferroni correction using the dunn.test package (98,99).

To statistically evaluate the clustering of samples by phage taxonomy, we used permutational multivariate analysis of variance (PERMANOVA) (100), implemented via adonis2 in the vegan package, at the phage family, genus, and species levels. Post-hoc pairwise PERMANOVA tests were performed using the pairwise.adonis function from the pairwiseAdonis package (101). Results of PERMANOVA are shown in **Tables S3-5.**

### Examination of differences in *Slopekvirus* long tail fibers

To understand the differences in the bacterial surface receptor genes for the *Slopekviruses* we examined the long-tail fiber protein that is required for phage adsorption (61). To build an MSA phylogenetic tree to look at where the phage tail fibers clustered based on the bacterial surface gene requirements, we input the AA sequences into NGPhylogeny. fr PhyML/OneClick advanced workflow using the standard settings and adding bootstrap calculations. Tree used in **Fig. 5A** was visualized with iTOL (102). The MSA alignment used in **Fig. 5B** was also exported from the workflow and visualized using Geneious Prime 2020.05.

## Supporting information

**S1 Figure.** Minimal impact of the *Klebsiella* ΔdsbA mutation on phage infectivity. (A) Efficiency of plating (EOP) for six phages with RB-TnSeq hits in ΔdsbA, measured on wild-type (WT, blue) and ΔdsbA (red) strains. (B) Difference in area under the curve (AUC) of bacterial growth (OD) between uninfected and phage-infected cultures in WT and ΔdsbA backgrounds. AUC was calculated in GraphPad, and to normalize for inherent growth differences between WT and ΔdsbA, the AUC of non-infected controls was subtracted from that of each phage-infected culture. Statistical significance comparing phage infection of WT bacteria to ΔdsbA was done using a two-tailed t-test in GraphPad Prism. P values are denoted as follows: P < 0.05 (*), P < 0.01 (**), and P < 0.001 (***).

**S2 Figure. STRING protein-protein interaction network of *Klebsiella* sp. M5al genes with high fitness scores from RB-TnSeq scores**. Nodes are colored according to STRING local network clusters: LPS biosynthesis (red), enterobacterial common antigen biosynthesis (green), O-antigen biosynthesis (yellow), and cellulose biosynthesis (blue). White nodes represent genes that do not belong to a local network cluster or KEGG pathway. Edge thickness indicates the confidence score of the predicted protein-protein interactions based on STRING’s scoring system with a minimum confidence score of 0.7.

**S1 Table. RB-TnSeq data used to generate figure 2**.

**S2 Table. Raw RB-TnSeq data.**

**S3-5 Table. Permanova of RB-TnSeq data at family, genus, and species level.**

**S6 Table. Proteins predicted to have TAT signaling domains in Klebsiella sp. M5al and phages.**

**S7 Table. Literature on bacterial genes previously found in phage infection.** Functions and references to bacterial genes that have been found in previous phage infection across different hosts and phages.

**S8 Table. Phages and bacteria used in the study.**

**S9 Table. Reasonings behind additional replicates for RB-TnSeq assays.**

**S10 Table. list of RB-TnSeq hits with significant, high fitness scores.**

**S11 Table. List of high fitness genes removed from RB-TnSeq assay and reasonings for removal.**

**S12 Table. List of genes identified in RB-TnSeq with hypothetical or unknown pathway functions.** We ran each of these genes through sequence similarity, hidden markov model profiles, domain searches, structural predictions, and literature search.

**S13 Table. Operon prediction for the *Klebsiella* sp. M5al genome.**

**S14 Table. gRNAs used for generating single gene Kos.**

**S15 Table. primers used to insert the KO gene back into Klebsiella sp. M5al.**

**S16 Table data used to generate the gephi graph examining the phage-bacterial gene interactions.**

## Notes

### Competing Interest Statement

The authors have declared no competing interest.

### Summary of Updates

This revised manuscript substantially expands both the experimental validation and conceptual framing of our RB-TnSeq based analysis of phage host interactions in Klebsiella pneumoniae. The study now presents the first systematic, high throughput comparative map of host factor requirements across 25 phages spanning five families, significantly extending the limited prior knowledge of phage receptors in this genus. We have strengthened the manuscript through the addition of extensive gene-specific validation experiments. Individual, sequence confirmed knockouts with complementation were generated for ten genes, including LPS outer core biosynthesis genes (wabI, wabN, wabH), the porin ompK36, the fatty acid transporter fadL, the cofactor biosynthesis gene pqqU, and additional host factors such as RS24775 and dsbA. These experiments confirm both known and newly identified determinants of phage infection, including several factors not previously associated with phage biology in any bacterial system. Validation assays include efficiency of plating, adsorption assays, and liquid growth inhibition experiments, substantially reinforcing the biological conclusions. We also clarify the interpretation and strengths of the RB-TnSeq approach. Gene-level fitness scores reflect concordant enrichment across multiple independent transposon insertions, providing internal replication and a robust measure of gene contribution to phage infection. Consistent with prior work, we present these results as high-confidence functional associations, supported by both internal reproducibility and targeted experimental validation. The manuscript has been reframed to emphasize its role as a high-throughput receptor mapping resource. The resulting dataset links diverse phages to host structures spanning LPS biosynthesis, outer membrane porins, transport systems, cofactor biosynthesis pathways, and regulatory networks. This resource provides a foundation for connecting phage genomic diversity to functional host interaction landscapes. We further provide a detailed rationale for the use of an acapsular host system. Capsule loss is the predominant mechanism of phage resistance in K. pneumoniae, and the receptor landscape characterized here reflects the secondary structures exposed in clinically relevant, phage resistant populations. As such, this work defines receptor targets of direct importance for therapeutic phage design. Finally, we refine terminology throughout to distinguish validated receptors, contributing host factors, and high-confidence candidates, and we highlight a key systems level finding: phage genus is a stronger predictor of host dependency profiles than phage family. Together, these revisions enhance the rigor, clarity, and translational relevance of the study while establishing a broadly useful community resource.

### Funding

- Department of Energy and Environment: #DE-SC0020173, #DE-SC0023307

