## Supplementary figures and images for "Cross-family and phage-specific gene requirements for Klebsiella infection revealed by scalable RB-TnSeq genetic screens"

### Supplemental Figure 1

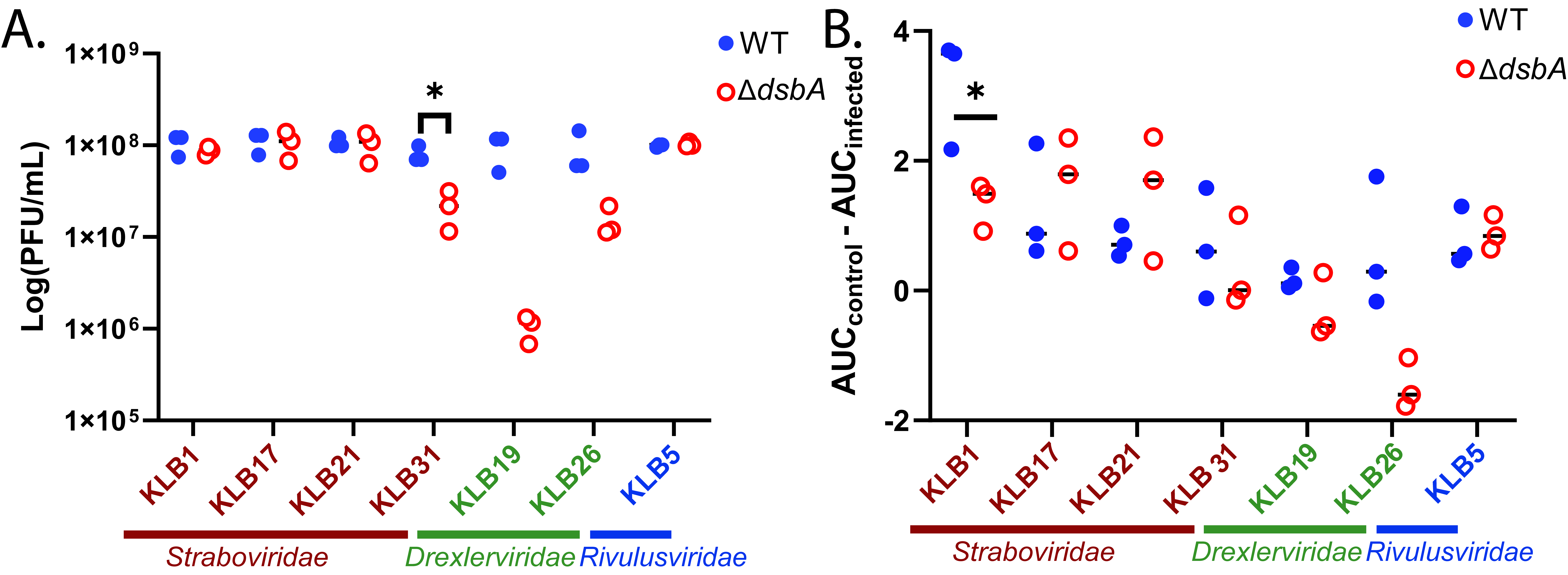

### Supplemental Figure 2

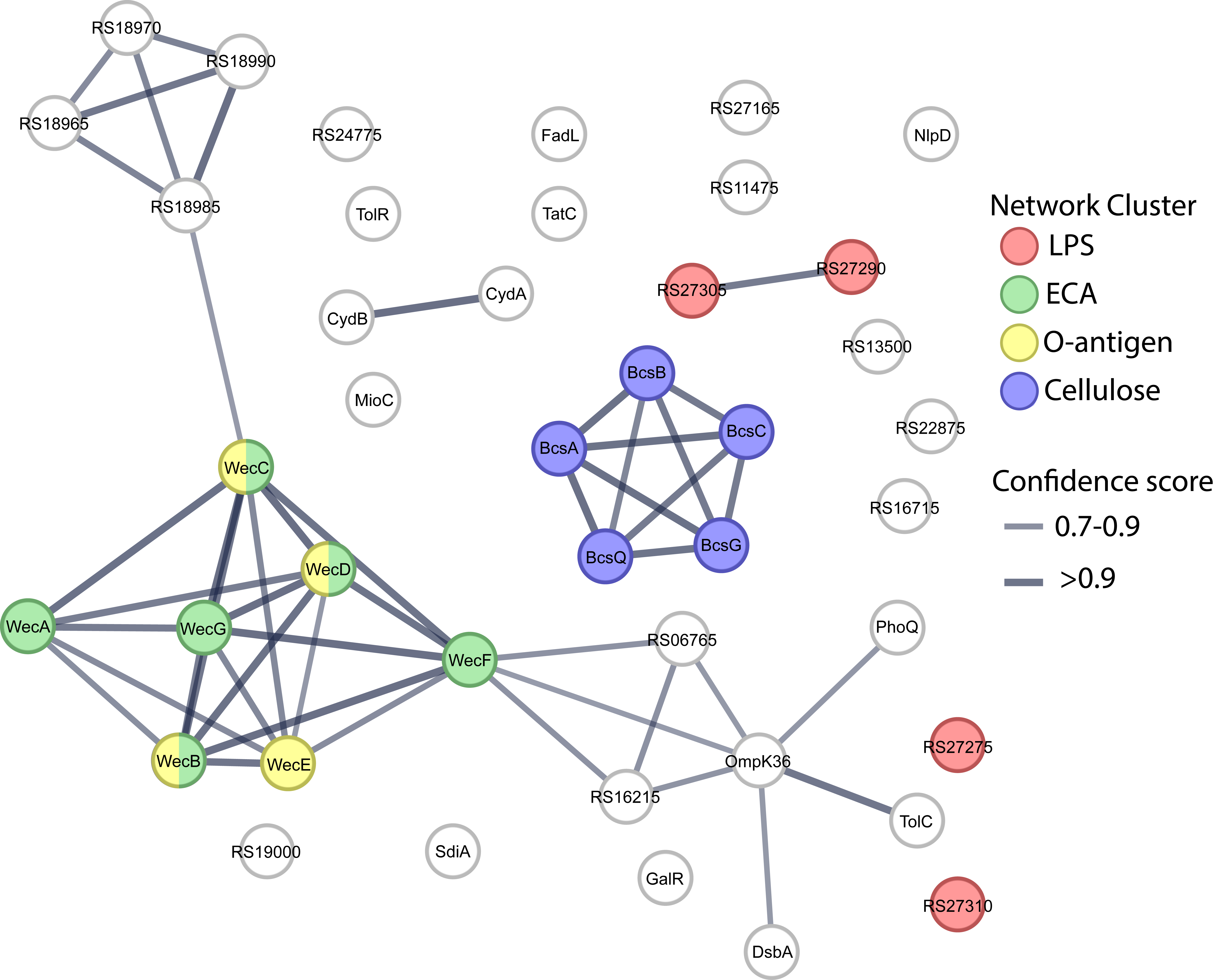
